# Plasmid architecture determines the stability of inverted terminal repeats in adeno-associated virus vectors

**DOI:** 10.64898/2026.09.24.753884

**Authors:** Vinod Jaskula-Ranga, Sheri Li, Aditya Pandya, Fred Bunz

## Abstract

Recombinant vectors derived from adeno-associated viruses (rAAVs) are a mainstay of human gene therapy. rAAVs are produced from plasmids containing transgene cassettes flanked by inverted terminal repeats (ITRs), which form structured DNA elements that stabilize the ends of the single-stranded viral genome and are the only viral sequences required in cis for genome packaging. For decades, it has been recognized that propagation of ITR-containing plasmids can result in deletions and other mutations, prompting the use of specialized bacterial strains, modified growth conditions, and truncated or altered ITRs. Despite these practices, ITR instability remains a persistent source of plasmid heterogeneity. To identify determinants of ITR stability, we evaluated ITR integrity in one of the original cloned AAV2 genome isolates, a reconstructed AAV2 plasmid, and a synthetic rAAV vector containing full-length native AAV2 ITRs. We established a quantitative bioinformatic workflow for analyzing ITR-containing plasmids and virus preparations from raw Oxford Nanopore sequencing data. These experiments showed that ITRs were highly stable during short-term culture, whereas prolonged culture revealed strong positional effects, with preferential loss or mutation of the ITR nearest the plasmid origin of replication. Consistent with this model, a survey of 7,041 sequence-verifiable AAV plasmids from the Addgene repository identified a widely disseminated 11-bp ITR deletion in 4,773 plasmids; among analyzable two-ITR plasmids, this deletion was located in the origin-proximal ITR in 95.3% of cases. Guided by these findings, we constructed a novel rAAV entry vector with stable full-length native AAV2 ITRs that enabled efficient packaging of a 4,750-bp all-in-one CRISPR-Cas9 cassette. Finally, we developed a cell-based strategy to compare the effects of ITR mutations on rAAV genome integration, providing preliminary evidence that ITR sequence variation can influence integration outcomes. Together, these findings show that ITR instability is a preventable, position-dependent property of plasmid architecture and identify ITR integrity as an important variable in rAAV vector design and quality control.

## Introduction

Recombinant AAVs derived from adeno-associated virus serotype 2 (AAV2) have been extensively employed as vectors for gene therapy (1,2). Unmodified AAV2 consists of a 4.7 kb single-stranded DNA genome that is delivered by a ∼20 nm icosahedral capsid. The native genome encodes two major open reading frames flanked by inverted terminal repeats (ITRs). These 145 nt elements form protective T-shaped duplex structures that prime second-strand DNA synthesis and serve as packaging signals during genome encapsidation.

The successful cloning of the AAV2 viral genome more than 40 years ago enabled the rescue of infectious virus from a recombinant plasmid (3,4). The ITRs are the only viral elements required for packaging in *cis*; the remaining viral components can be replaced by transgene cassettes. These “gutless” recombinant genomes can be packaged by introducing the essential AAV and adenovirus genes in *trans*. In practice, rAAV packaging is usually performed by co-transfection of a *cis* plasmid vector containing AAV2 ITRs with plasmids that express the complementary AAV and adenoviral elements. To exploit the varied tropism of individual AAV serotypes, researchers have successfully packaged AAV2 ITR-flanked transgenes in a variety of naturally occurring and engineered capsids (5), an approach known as pseudotyping. For research and therapeutic use, rAAV vectors are valued for their ability to infect dividing and non-dividing cells, their episomal persistence, and their favorable safety profile (1).

Despite many technological advances, persistent challenges remain in large-scale rAAV manufacturing (6–8). Homogeneity and consistency are particularly important considerations for clinical applications.. Despite ongoing efforts to standardize the reagents and methods used for rAAV production, heterogeneity within individual virus preparations and variation between preparations continue to be problematic. Viral stocks typically consist of a variable mixture of functional virions and defective particles. The fraction of non-functional rAAV particles can vary significantly between preparations and typically includes empty capsids and virions that contain single-stranded DNA genomes with deletions or contaminating sequences (9). For clinical use, defective particles can reduce potency and increase treatment-associated toxicities (10).

One important contaminant in rAAV preparations is the backbone of the *cis* plasmid. Non-functional virions can be generated via a phenomenon known as reverse packaging, wherein the plasmid backbone is encapsidated instead of the desired cargo insert, creating what are referred to as snapback genomes that have a single ITR. Snapback genomes have been shown to arise from ITR-like sequences outside the flanked transgene that in effect create cryptic resolution sites for the replicase complex (11).

Another potential source of heterogeneity within rAAV preparations is heterogeneity in the rAAV *cis* plasmid stocks used to produce them. There is a well-documented tendency for AAV ITRs propagated in standard *cis* plasmids to be unstable (12). More broadly, highly structured viral sequences and repetitive elements of many kinds frequently accumulate deletions and other types of mutation when propagated in recombinant plasmid vectors (13). ITR instability is generally believed to be a consequence of their palindromic structure and high GC content, which make them particularly prone to hairpin formation and replication stalls. ITR mutations can compromise rAAV packaging efficiency, yield, and *in vivo* efficacy (14). To minimize ITR defects, manufacturers commonly use a variety of modified culture conditions, such as growth at reduced temperature, and use specialized strains. Despite these measures, the stable propagation of repetitive elements remains problematic.

A retrospective look at the earliest AAV2 rescue experiments suggests that the two ITRs differ in their propensity for mutation when propagated in plasmid vectors (15). The relative positions of the two ITRs with respect to the origin of replication has been suggested to contribute to ITR instability (16). In this study, we established a workflow that permitted detailed examination of how plasmid configuration influences ITR instability in founding AAV2 isolates and synthetic rAAV vectors. In both low-copy and high-copy vectors, ITR stability was found to be related to the position of the ITRs with respect to the plasmid origin of replication. A simple vector modification eliminated this positional effect and restored plasmid stability.

## Materials and Methods

### Plasmids

The AAV2 plasmids pSM620 and psub201 were acquired from the National Gene Vector Biorepository (NGVB). The plasmids pAAV-RC and pHelper were originally purchased from Stratagene, now Agilent Technologies (La Jolla, CA), as components of their AAV-Helper Free kit.

Plasmid DNAs were propagated in the *E. coli* DH5α derivative NEB5 (New England Biolabs, Ipswich, MA). This strain has the genotype: *fhuA2::IS2* Δ*(mmuP-mhpD)169* Δ*phoA8 glnX44 80d[*Δ*lacZ58(M15)] rfbD1 gyrA96 luxS11 recA1 endA1 rphWT thiE1 hsdR17*. Chemically competent bacteria were transformed by heat shock and cultured in Luria-Bertani (LB) broth or on LB agar plates. Depending on the selectable marker, the conditions for bacterial selection were 100 µg/ml carbenicillin, 50 µg/ml kanamycin or 20 µg/ml doxycycline.

Two 145-bp AAV2 ITRs were synthesized in separate plasmids by Genscript, Inc. (Piscataway, NJ), both in the flop orientation. The left ITR was synthesized in pUC57-KAN; the right ITR was synthesized with a 1,577-bp green fluorescent protein (GFP) expression cassette in pUC57-AMP. The right ITR-GFP cassette was excised and inserted into compatible sites in the left ITR plasmid to create the plasmid pAAVcis. This plasmid was further modified by separating the *ori* from the right ITR with a stuffer fragment. A 1,320-bp segment of pBR322, which contains the *Tet^r^* open reading frame, was amplified with the forward primer 5’-AACACATGTTCTCATGTTTGACAGCTTATC-3’, and the reverse primer 5’-accACATGTCTTGGAGTGGTGAATCCG-3’. The amplified DNA was gel-purified and the ends were digested with PciI for ligation into the unique PciI restriction site in pAAVcis. The resulting pAAV2ST plasmid was deposited with Addgene (plasmid #239400; http://n2t.net/addgene:239400; RRID:Addgene_239400).

The pAAV2ST-Pcsk9sg1 plasmid was constructed by first introducing the Pcsk1sg1 protospacer 5’-CACCGCAGCCACGCAGAGCA-3’ (17) into the plasmid pX602-AAV-TBG::NLS-SaCas9-NLS-HA-OLLAS-bGHpA;U6::BsaI-sgRNA (a gift from Dr. Feng Zhang, Addgene Plasmid #61593) and then amplifying a 4,428-bp NotI-NotI insert with the forward primer 5’-CCTGCGGCCGCTCTAGACTCGAGGGGCTGGAAG-3’ and the reverse primer 5’-CCTGCGGCCGCGAGGGCCTATTTCCCATGATTC-3’. The resulting fragment was cloned into NotI-digested pAAV2ST.

The pBR-AAV2(-) and pBR-AAV2(+) plasmids were constructed in two stages. First, a pAAVcis fragment containing both ITRs and the GFP transgene was inserted into the NheI and BamHI sites of pBR322. Next, the AAV2 genomic sequences between the two ITRs in pSM620 were amplified with Platinum SuperFi II Master Mix (Thermo Scientific) using the primers 5’-ATCACTAGGGGTTCCTGGAGGGGTGGAGTCGTGACG-3’ and 5’-ATCACTAGGGGTTCCTTGTAGTTAATGATTAACCCG-3’. The resulting insert was seamlessly cloned into the NotI-digested intermediate vector with the NEBuilder kit (New England Biolabs, Ipswich, MA). The annotated sequences for the following plasmids have been deposited in GenBank: pSM620 (accession **PZ196606**), pAAV2ST (accession **PX505299**), pAAV2ST-Pcsk9sg1 (accession **PX505298**).

### Viral packaging and analysis

The pAAV2ST-Pcsk9sg1 plasmid was amplified at an industrial scale and subsequently packaged into AAV8 capsids by VectorBuilder, Inc. (Chicago, IL). The purified preparation was first assessed by mass photometry by SignaGen, Inc (Frederick, MD). For analysis by transmission electron microscopy (TEM), particles were directly applied to a grid and negatively stained with 1% phosphotungstic acid. Sample preparation and imaging were performed at the Johns Hopkins University School of Medicine Microscope Facility.

### DNA sequencing

The ITR regions of AAV2 plasmids were sequenced by Genewiz/Azenta (South Plainfield, NJ) using a proprietary Sanger sequencing protocol optimized for hairpin structures. The sequencing primers were: Right ITR: 5’-ACTTTGGTCTCTGCGTATTTCT-3’; Left ITR: 5’-AACACTCACGTGACCTCTAATAC-3’.

Oxford Nanopore Technologies (ONT, Oxford Science Park, UK) long-read sequencing of the plasmid constructs was performed by Genewiz/Azenta. Briefly, plasmid samples were subjected to library preparation optimized for circular DNA templates and sequenced on ONT flow cells using standard ligation chemistry. Raw nanopore reads were basecalled by the provider using the Oxford Nanopore dna_r10.4.1_e8.2_400bps_sup-v4.1.0 model (high accuracy SUP model for R10.4.1 chemistry).

Per-position basecall heterogeneity across plasmid sequences was assessed by a custom Python pipeline applied directly to raw ONT FASTQ reads. Reads were aligned to circular reference sequences using a seed-chain algorithm with banded Needleman-Wunsch gap filling. Briefly, a k-mer index (k = 12, stride = 3) was constructed from the reference sequence, and for each read, strand orientation was determined by k-mer hit density. Alignment offset was estimated by diagonal voting across k-mer hits. Reads spanning the reference linearization point were detected by identification of a second dominant diagonal offset by approximately one plasmid length. Aligned bases were recorded at each reference position, and inter-anchor gaps were resolved by banded Needleman-Wunsch alignment (half-bandwidth = 12). Read termini were aligned by the same procedure over a maximum of 80 bp. For each reference position, coverage and per-base counts (A, T, G, and C) were tabulated, and a deviation score was computed as the fraction of basecalls inconsistent with the reference base. Positions with deviation ≥ 10% at coverage ≥ 50× were flagged for further inspection.

All analyses were performed using Python 3.12 with NumPy, pandas, and matplotlib. No external alignment software was used. Reference sequences were linearized at a position distal from both ITRs (> 2 kb) to prevent wrap-around alignment artifacts at palindromic sequence boundaries.

Oxford Nanopore sequencing reads from encapsidated AAV viral genomes were analyzed using a custom quality control pipeline (aav_genome_qc.py). Inverted terminal repeats (ITRs) were automatically detected from the full-length cis plasmid reference sequence based on palindromic structure and serotype-specific motifs. Reads were aligned to the cis plasmid reference using minimap2 through mappy with the “map-ont” preset. Contamination analysis was performed by competitive mapping to helper plasmid, rep-cap plasmid, and host (*E. coli*) reference sequences, with reads assigned to the reference yielding the highest mapping quality score (minimum MAPQ ≥ 20). Transgene-mapping reads were structurally classified into full-length single-stranded AAV (ssAAV), full-length self-complementary AAV (scAAV), partial ssAAV (5’ or 3’ incomplete genome fragments), partial scAAV (snapback genomes), or reverse-packaging products based on alignment coordinates relative to ITR boundaries. Per-position coverage and strand-specific coverage were calculated across the transgene insert region, with base quality filtering (minimum Q ≥ 10). ITR coverage statistics were computed separately for both terminal repeats. Truncation hotspots were identified from alignment start and end coordinates of transgene-mapping reads. All results were exported as CSV files and visualized as graphs from Excel.

All FASTQ files are available in the Sequence Read Archive (**PRJNA1439403** released 2026-03-19). Analysis code is available at https://github.com/fredbunz-lgtm/aav-itr-pileup (DOI: 10.5281/zenodo.19186494.)

### Survey of the Addgene plasmid repository

Addgene plasmid sequence data were accessed through the Addgene Developers API using token-based authentication. Because catalog search access was restricted, the bulk plasmid-with-sequences download was used for sequence analysis. The resulting JSON file was streamed to avoid loading the full dataset into memory. A metadata-based screen was first used to identify candidate AAV-related plasmids. Candidate plasmids were then scanned for AAV ITR sequences using a local alignment-based ITR detection workflow. Sequence-level ITR calls were collapsed to plasmid-level summaries using representative sequence records to avoid counting multiple sequence files from the same plasmid as independent biological ITRs. When multiple sequence records were available for a plasmid, representative records for ITR structural classification were selected based on the presence and number of detectable ITRs.

ITR calls were classified by alignment to AAV2 ITR reference sequences representing both flip and flop configurations and both deposited strand orientations. For each ITR call, the best-matching reference was used to infer the flip/flop configuration. ITR structural classes were assigned from the aligned reference span, aligned plasmid span, and Δ11 junction status. ITRs were classified as full-length 145-bp ITRs, 130-bp truncated ITRs, 119-bp Δ11 ITRs, 119-like ITRs lacking the exact Δ11 motif, or other altered/truncated ITRs. Plasmids containing two inferred ITRs were then summarized by paired ITR architecture. pAAV2ST (Addgene plasmid 239400), served as a positive control for detection of two full-length 145-bp ITRs.

The prevalence of the Δ11 mutation was assessed by searching for the 34-bp sequence corresponding to the 11-bp deletion, in both orientations. Circular sequences were handled by appending the first 33 bases of each sequence to its end, allowing detection of motif instances spanning the sequence linearization point without duplicating internal motif counts. A known Δ11-containing control sequence was used to verify that the exact-motif workflow reported a single expected motif instance.

For origin-proximity analysis, bacterial replication origin coordinates were obtained from GenBank **rep_origin** feature annotations. Bulk sequence-derived ITR and Δ11 coordinates were compared with GenBank sequence coordinates after confirming sequence-length, sequence-identity, and Δ11-position concordance between representative bulk and GenBank records. Plasmids were included in the primary origin-proximity analysis only if the representative sequence contained exactly two inferred ITRs and exactly one exact Δ11 motif. The circular distance from each ITR midpoint to the bacterial origin midpoint was calculated, and the Δ11-containing ITR was classified according to whether it was the closer or farther ITR relative to the annotated plasmid ori element.

### Genomic integration assay

The gene targeting vector pSEPTp53 (18) contains a *TP53* knockout cassette consisting of two ∼1-kb homology arms that flank a synthetic exon promoter trap. This cassette had been cloned into the NotI sites of the AAV entry vector pAAV-MCS (Agilent). The pSEPTp53 plasmid was digested with NotI, and the dephosphorylated insert was re-inserted into NotI-digested pAAV-MCS in either orientation. The same insert fragment was directly cloned into the NotI sites of the pAAVcis plasmid. The three resulting plasmids were packaged into AAV2 capsids with the AAV Helper-Free System (Agilent), as described (18). Unpurified virus stocks were titered by qPCR and added to HCT116 and hTERT-RPE1 cells growing in sub-confluent monolayer cultures at a multiplicity of infection of 10. Three days after infection, cells were detached with trypsin-EDTA and plated at limiting dilution to 96-well cell culture plates (Corning) in medium supplemented with 0.4 mg/ml Geneticin (ThermoFisher Scientific). Between 60 and 110 individual colonies were scored from each set of plates and genomic DNA was directly extracted with Lyse-N-Go PCR reagent (Pierce Biotechnology, Rockford, IL). A PCR screen for homologous integration into the *TP53* locus was performed with Platinum Taq (ThermoFisher Scientific) according to manufacturer’s recommendations using the primers: p53R, 5’-TGTGTGTGACTGCTTGTAG-3’; NeoFor, 5’-TCTGGATTCATCGACTGTGG-3’. Samples that generated a 1.8-kb positive signal were confirmed by a second PCR with primers: p53F, 5’-CTGGGAGAAGGTGCGATGAT-3’; NeoRev, 5’-GTTGTGCCCAGTCATAGCCG-3’, which generated a 2.2-kb band from homologous integration events.

## Results

### Characterization of ITRs in the original AAV2 plasmids

The original cloning of the native AAV2 genome required the conversion of a native single-stranded genome into a double-stranded plasmid insert. In this form, the ITRs can either self-anneal to assume a cruciform structure or form a linear duplex with the complementary DNA strand (**Fig 1a**). To determine how plasmid elements outside the AAV2 insert might influence this structural equilibrium or otherwise affect ITR stability, we studied the plasmid pSM620, which was among the first cloned AAV2 isolates (3). We also examined the pSM620 derivative psub201 (19), which includes XbaI restriction sites that permit the entire region between the two ITRs to be replaced with transgenic cargo (**Fig 1b**). The psub201 plasmid was the forerunner of many of the gutless entry vectors that are now widely used for rAAV production.

**Fig 1.**
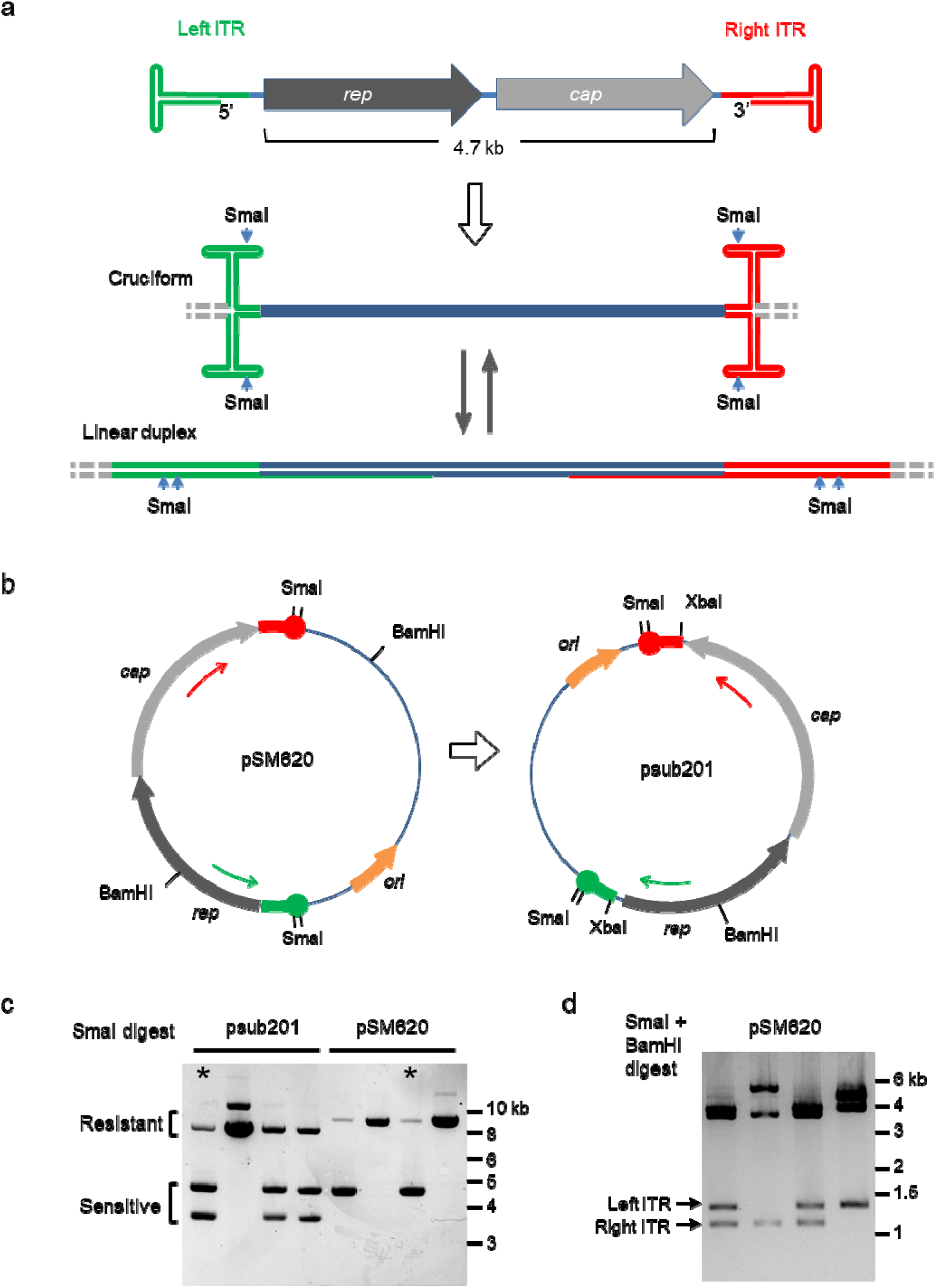
The original AAV2 plasmids. (a) The single-strand DNA genome of AAV2 consists of two transcriptional units, *rep* and *cap*, flanked by ITRs. In a double-stand DNA context, each ITR can form a linear or cruciform structure. (b) The original AAV2 isolate pSM620 was used to derive psub201. The wild-type 145-bp ITRs were truncated to 130-bp during the cloning process. The XbaI sites flanking the *rep* and *cap* genes permitted the replacement of the endogenous genomic elements with transgene cassettes. The relative positions of the primers used for Sanger sequencing are indicated by arrows. (c) Digestion with SmaI reveals ITR heterogeneity among individual clones isolated from the reference stocks of pSM620 and psub201. (d) The pSM620 clones analyzed in panel c were digested with both SmaI and BamHI to identify the deleted ITR.

Reference samples of pSM620 and psub201 DNAs were amplified by bacterial transformation and single-colony plating. Four colonies were randomly selected from each plate and grown overnight at small scale. The extracted DNAs were first analyzed by restriction digest. Each ITR contains two recognition sites for the restriction endonuclease SmaI (**Fig 1a**), which is routinely used to assess ITR integrity. Three of the eight plasmid isolates contained an ITR that was SmaI-resistant (**Fig 1c**). A double digest of pSM620 subclones revealed that each of the two ITR-defective clones had retained a different ITR (**Fig 1d**).

Isolates of pSM620 and psub201 that could be fully digested with SmaI were next assessed by Sanger sequencing. The secondary structure of AAV ITRs has long been a barrier to ITR sequencing, but refined techniques now permit the reliable extension of sequencing primers through GC-rich hairpins and other highly structured sequence motifs (20,21). Using primers that annealed within the AAV2 genome (**Fig 1b**), we confirmed that SmaI sites were present in both ITRs in each of the two plasmids. However, the sequence quality differed between ITRs in each plasmid, even though the same primers were used in both. The pSM620 template yielded high-quality sequence across the right ITR, while basecalling confidence was consistently lower across the left ITR (**Fig 2a**). The opposite was true for the psub201 plasmid; in this case the right ITR yielded sequence data that were relatively poor in quality, but nonetheless sufficient to reveal a 21-bp deletion (**Fig 2b**).

**Figure 2.**
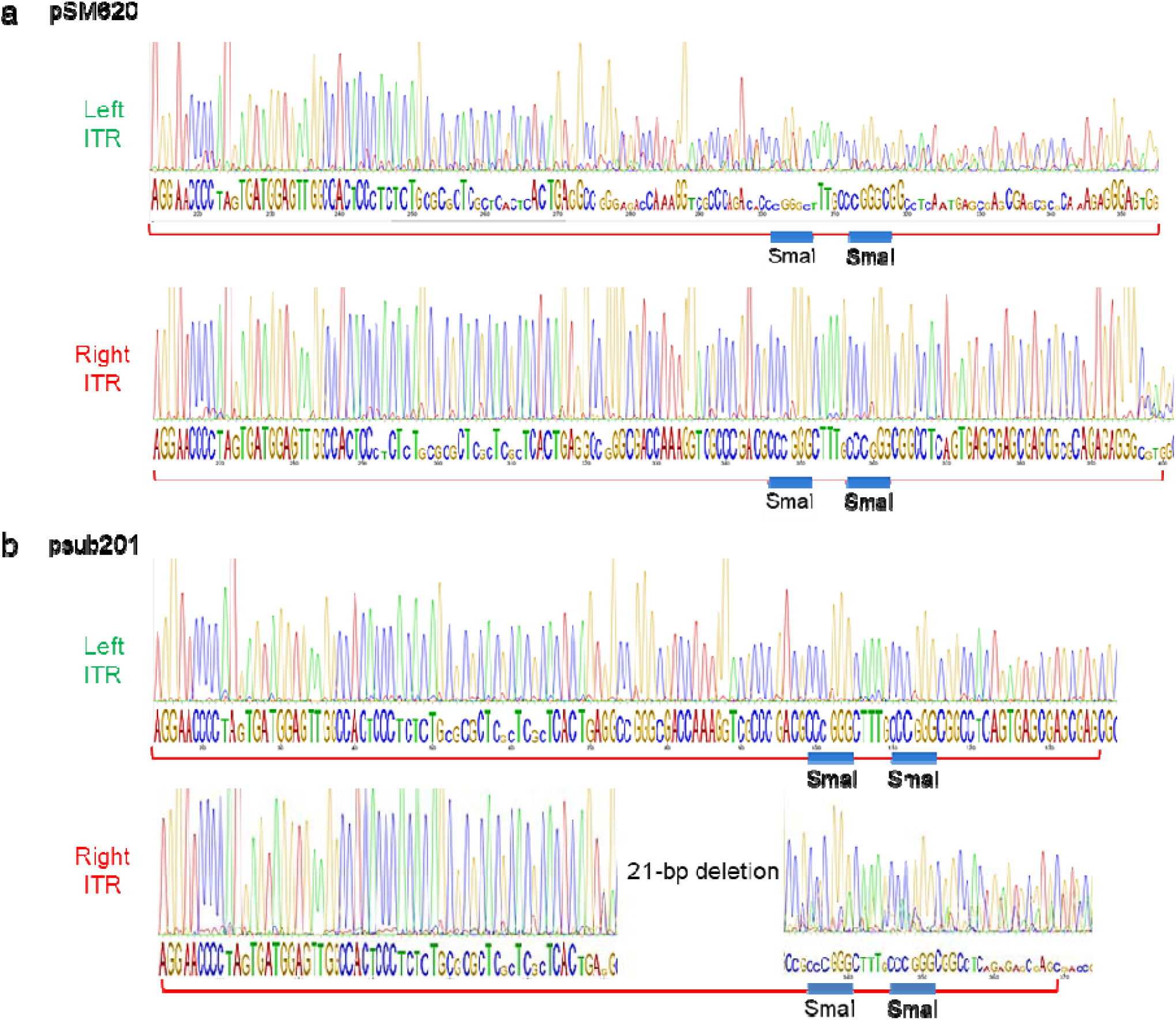
Asymmetric Sanger sequencing quality across ITRs. The left and right ITRs of the SmaI-sensitive (a) pSM620 and (b) psub201 subclones marked by asterisks in Fig 1c were sequenced by a modified Sanger method with the primers shown in Fig 1b. The relative confidence of the basecall at each position is indicated by letter height.

To deconvolute what appeared to be heterogeneous plasmid template sequences, the ITR elements in pSM620 were further characterized at the single-molecule level by long-read DNA sequencing. Nanopore reads above the minimum quality cutoff were homogeneous across both ITRs (**Fig 3a, b**). This analysis resolved a 143-bp left ITR 136-bp right ITR, both in the flop orientation. Two bases were missing from the 5’ end of the left ITR and nine bases were missing from the 3’ end of the right ITR. A small proportion of the reads contained small indels, occurring at the highest frequency at the 5’ end of the left ITR (**Fig 3a**). In general, the ITR regions exhibited the largest degree of heterogeneity, as evidenced by off-mode base calls in the pileup (**Fig 3c**).

**Figure 3.**
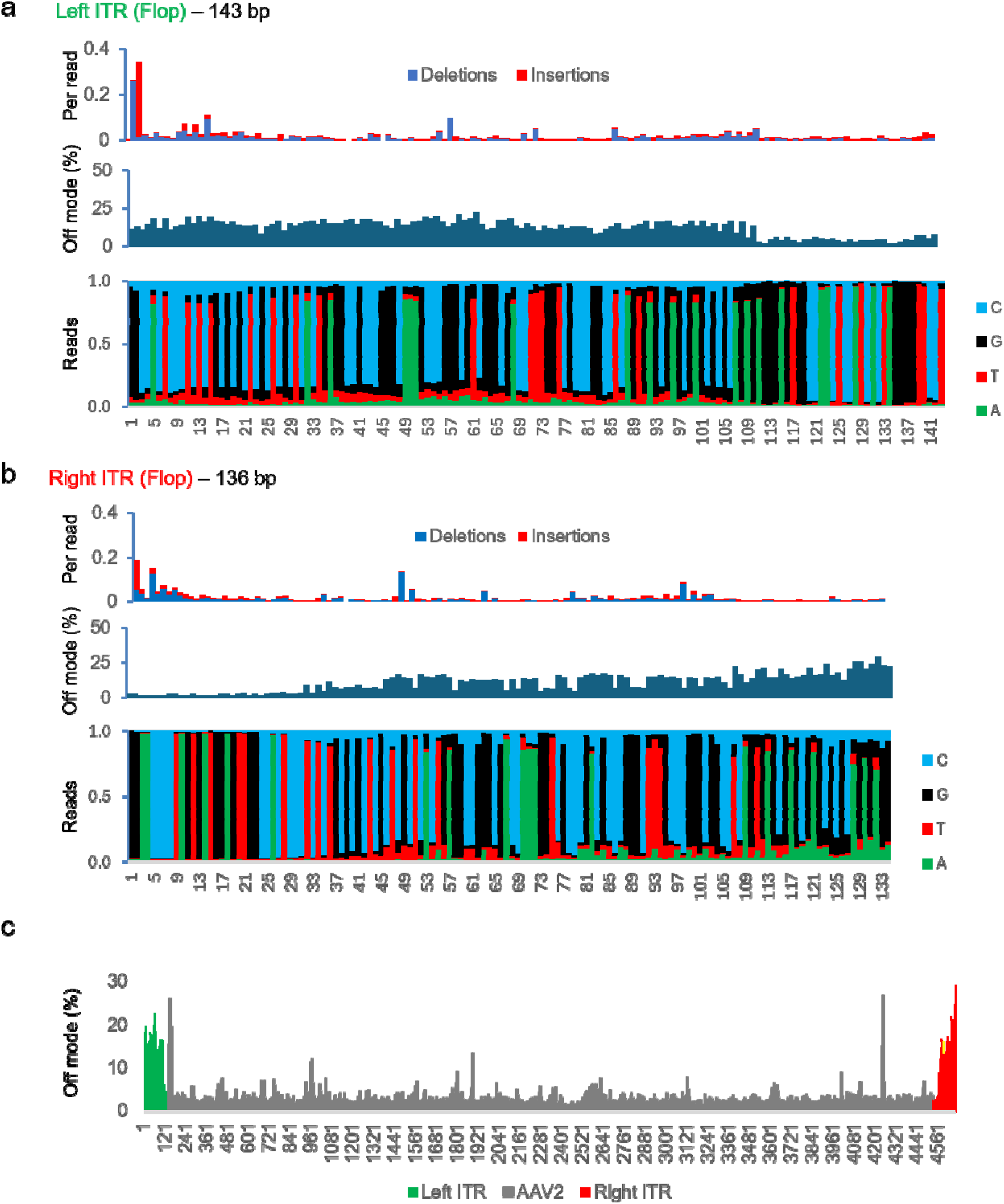
Nanopore sequencing across pSM620 ITR elements. Read pileup analysis across the left ITR (a) and the right ITR (b) is graphically illustrated. A stacked bar graph shows the proportion of bases at each position, as indicated on the horizontal axis. Identified insertions and deletions are represented as the fraction of all reads at each position. The off-mode plot represents the divergence of the reads from the reference sequence at each position. (c) The sequence divergence at each position across the AAV2 insert in pSM620.

### Reconfiguration of pSM620

As the asymmetric quality of the Sanger traces was not reflected in the underlying pSM620 template sequence, we investigated whether the poor Sanger sequencing quality from the left ITR might be related to structural constraints on primer extension. In both pSM620 and psub201, the affected ITR was adjacent to the plasmid origin of replication (**Fig 1b**). The pMB1 *ori* element, a ColE1-type replication origin, expresses two short RNAs that control plasmid copy number (22). We wondered whether proximity to the template elements for these highly structured RNAs might account for the discordant sequencing data.

The pSM620 plasmid was originally derived by inserting the purified AAV2 genome into the PstI site of pBR322 by a capture strategy that involved terminal homopolymer tailing (3). The insertion site in the parent pBR322 plasmid is separated from the pMB1 *ori* element by less than 500 bp (**Fig 4a**). To investigate the possible positional effects of the replication origin on adjacent ITRs, we constructed new plasmids by inserting an AAV2 cassette into a pBR322 vector digested with NheI and BamHI. The resulting plasmids pBR-AAV2(+) and pBR-AAV2(-) contained the entire AAV2 genome, including two 145-bp ITRs, in either orientation. The repositioned and fully restored ITRs were spaced 1457 bp or 2144 bp from the pMB1 origin (**Fig 4a**).

**Figure 4.**
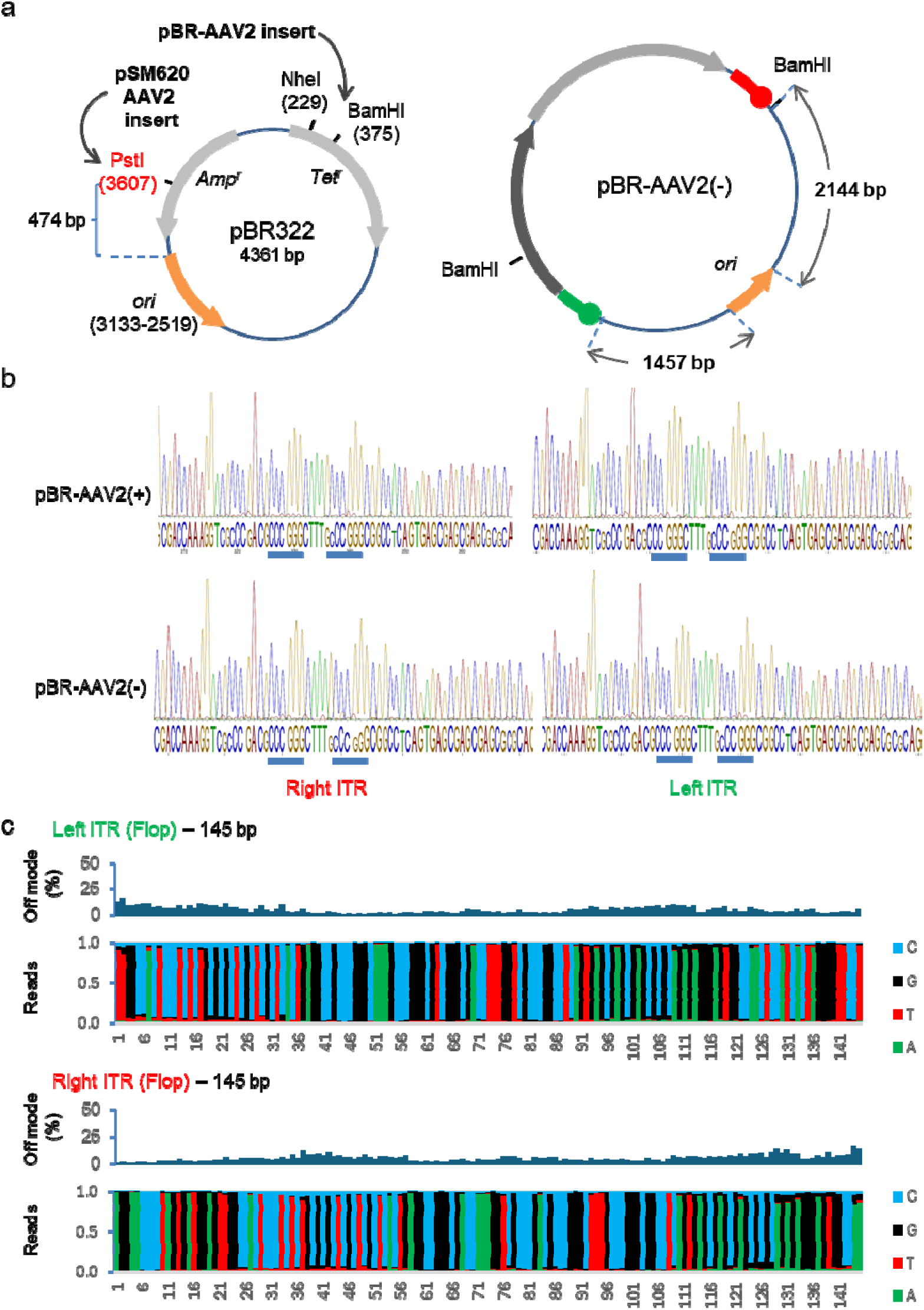
Revision of pSM620. (a) The original AAV2 isolate pSM620 was captured in the Pstl site of the cloning vector pBR322, thereby disrupting the gene for ampicillin resistance. To alter the positioning of the ITRs with respect to the *ori* element (orange arrow) the AAV2 genome was reconstructed as an NheI-BamHI insert, as shown. The resulting plasmid, pBR-AAV2(-) has the insert in the same orientation as pSM620, with respect to the direction of the *ori* element. A second plasmid, pBR-AAV2(+), features the AAV2 insert in the opposite orientation and is not shown. Numbers in parentheses refer to the nucleotide positions in pBR322. (b) Sanger sequencing was performed with the primers shown in Fig 1b. The confidence of the basecall at each position is indicated by letter height. SmaI recognition sites are underlined. (c) Nanopore sequencing pileup analysis across the left and right ITRs of pBR-AAV2(-).

The ITRs in pBR-AAV2(+) and pBR-AAV2(-) were assessed by Sanger sequencing with the same primers and conditions used for the prior analysis of pSM620 and psub201. Both ITRs were well-resolved in the Sanger traces, irrespective of the orientation of the AAV2 insert (**Fig 4b**). The ONT pileup analysis revealed low levels of heterogeneity across both ITRs (**Fig 4c**).

Next, we assessed ITR stability during extended culture. We compared pSM620 and pBR-AAV2(-), in which the AAV2 cassette was in the same directional orientation with respect to the replication origin (**Fig 4a**). Analysis by SmaI digestion showed that both plasmids remained structurally intact over a 72 h time course, with no apparent deletions or rearrangements (**Fig 5a**). Although all four recognition sites had been confirmed by DNA sequencing (**Fig 4b**), the SmaI enzyme consistently failed to digest the pSM620 plasmid to completion. Moreover, the extended culture of pSM620 appeared to cause a moderate increase in the relative proportion of SmaI-resistant DNA (**Fig 5a**). Long-read sequences of pSM620 DNAs isolated from the 72 h culture revealed the loss of 6-bp from the 5’ end of the left ITR but otherwise showed no evidence of expanded subclonal plasmid populations (**Fig 5b**).

**Figure 5.**
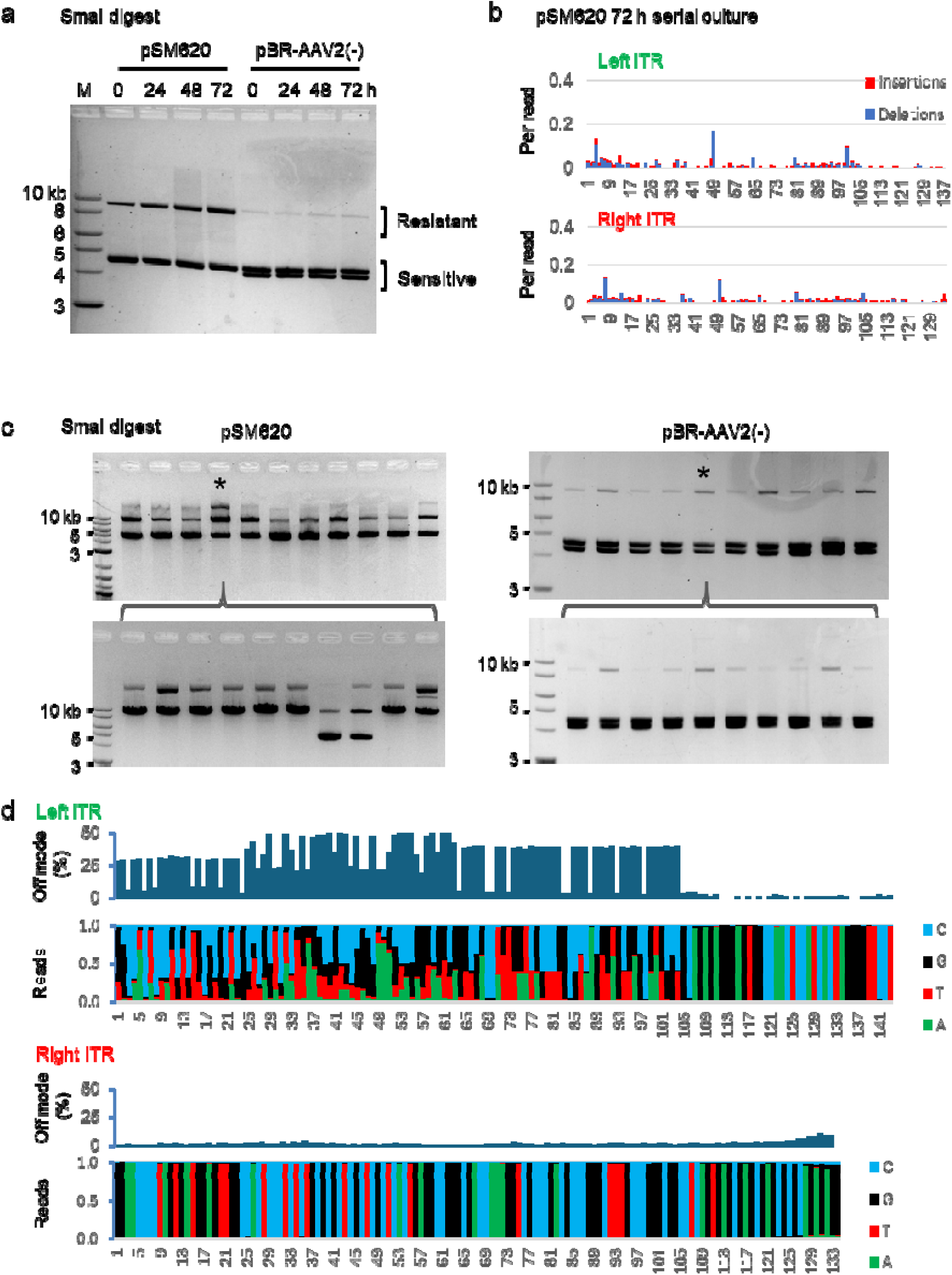
Stability of AAV2 plasmids during extended culture. (a) Liquid bacterial cultures were incubated continuously for 72 h. At 24-h intervals, an aliquot of each culture was serially diluted at a ratio of 1:1000 into fresh growth medium. DNAs extracted at each time point were digested with Smal. (b) Analysis of insertions and deletions across the left and right ITRs in pSM620 following 72 h of continuous culture. (c) The 72-h cultures were streaked on fresh plates, and 10 colonies were individually expanded in overnight cultures. Extracted plasmid DNAs were digested with SmaI. The cultures indicated by asterisks (*) were streaked out for a second round of subclonal analysis. (d) Nanopore sequencing pileup analysis across the left and right ITRs of pSM620 clone 4, as indicated in panel c.

To more deeply assess stability during extended culture, we used the 72 h DNA samples to transform bacteria and then individually expanded 10 random drug-resistant colonies. The extracted DNAs were assessed by SmaI digest, revealing variability among the subclones **(Fig 5c**). The pSM620 isolate with largest proportion of undigested DNA (clone 4) was further subcloned. The SmaI digestion pattern revealed a high rate of ITR deletion among these subclones. In contrast, subclones derived from pBR-AAV2(-), were relatively homogeneous, even following a second round of subcloning and expansion (**Fig 5c**). The heterogeneity in the clone 4 plasmid population was readily detectable by long-read sequencing (**Fig 5d**) and completely localized to the left *ori*-proximal ITR.

### Stability of 145-bp ITRs in a high-copy plasmid

The rAAV vectors commonly used today harbor a mutated pMB1 origin that permits plasmids to be maintained at a high copy number. To assess ITR stability in the context of a high-copy plasmid, we constructed a *cis* vector *de novo*. A 145-bp right ITR and a 145-bp left ITR were separately synthesized in the pUC57 backbone, both in the flop orientation to match the conformation of pSM620. These two plasmids were then used to assemble the *cis* plasmid, pAAVcis, in which a green fluorescent protein (GFP) expression cassette is flanked by two ITRs (**Fig 6a**).

**Figure 6.**
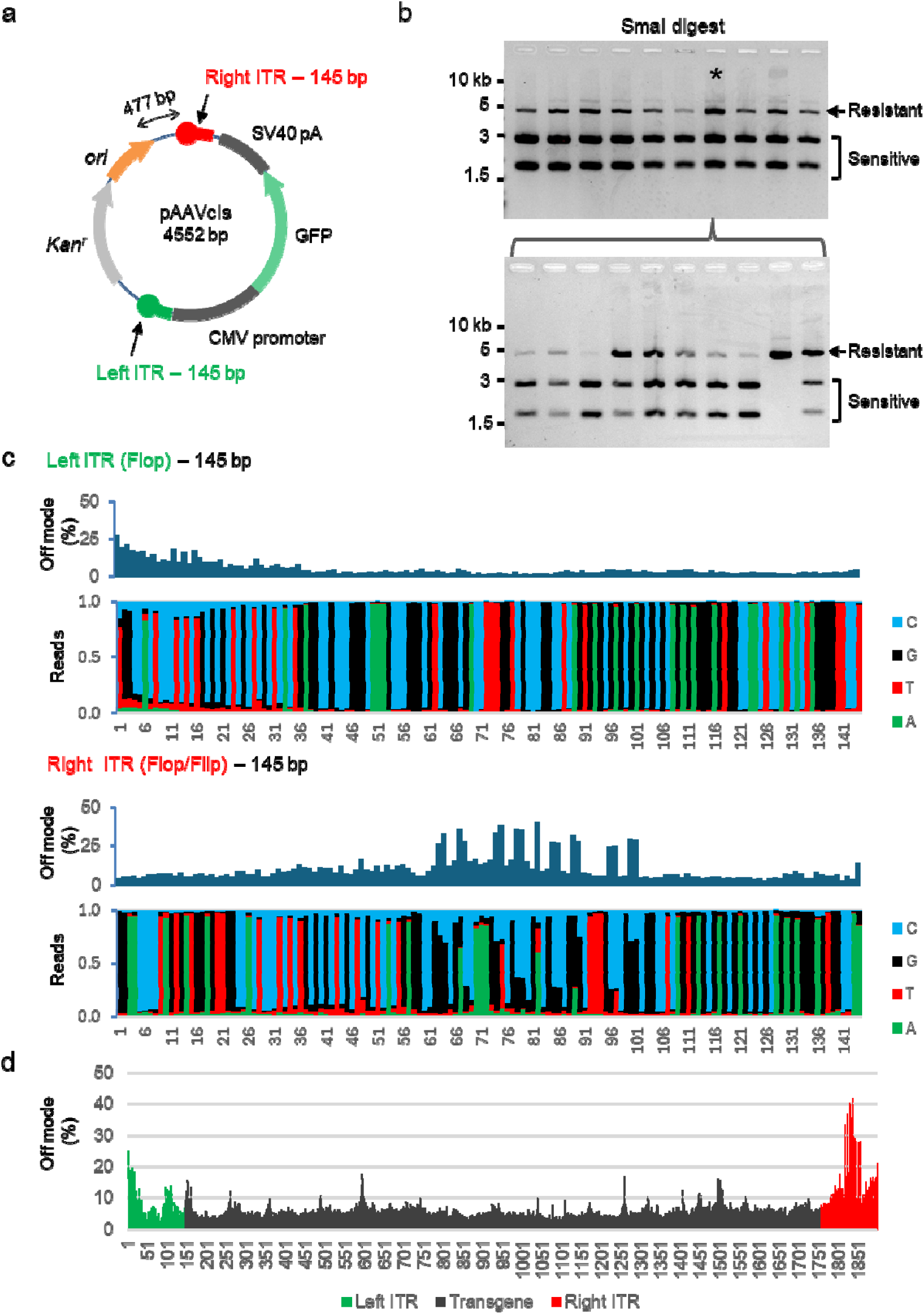
Analysis of a synthetic rAAV *cis* plasmid. (a) A GFP expression cassette flanked by 145-bp AAV2 ITRs was assembled in the high-copy cloning vector pUC57-KAN. (b) Following continuous culture for 72 h, 10 independent subclones were expanded and the extracted DNA was digested with Smal. The culture indicated by an asterisk (*, clone 7) was streaked on a fresh bacterial plate for a second round of subclonal analysis. (c) Nanopore sequencing analysis of the ITRs from pAAVcis clone 7. (d) Off-mode basecalls in the nanopore pileup data across the rAAV insert.

The pUC series of cloning vectors features a multiple cloning site that is positioned near the high-copy pMB1 origin. In pAAVcis, derived from pUC57-KAN, the plasmid origin is separated from the insertion site of the right ITR by 477 bp (**Fig 6a**). Using the iterative subcloning strategy employed for the analysis of the pBR322-based plasmids, we isolated pAAVcis clones with ITR deletions. To assess ITR stability and the homogeneity of the pAAVcis plasmid population, we plated transformed bacteria at limiting dilution and expanded 10 kanamycin-resistant subclones for further analysis (**Fig 6b**). Digestion with SmaI revealed apparent variation among the pAAVcis subclones, with differing proportions of SmaI-sensitive and -resistant DNAs.

A single clone (clone 7; **Fig 6b**) that appeared to yield a disproportionate level of undigested material was further assessed by nanopore sequencing. The reads across the left ITR were relatively homogeneous, but there was detectable sequence heterogeneity among the reads that mapped to the *ori*-adjacent right ITR (**Fig 6c**). Interestingly, the sequence analysis revealed a mixture of flip and flop isomers in the right ITR region. In contrast to the flip-flop configuration of pSM620, both ITRs in pAAVcis were created in the flop orientation and were therefore identical. Following extensive propagation, approximately 27% of the plasmid molecules harbored a distinct sequence variant within the inner loop of the *ori*-proximal right ITR, with each of the 17 informative positions in the inner loop indicating a minor population with flip orientation (**Fig. 6c**). In contrast, the origin-distal left ITR was homogenously in the flop orientation (< 1% flip at corresponding positions). These data suggest that a pure pAAVcis plasmid population diverged into a mixture of plasmids with both flip and flop ITR configurations appearing in the origin-proximal ITR. An analysis across the entire rAAV insert revealed that a distinct peak of heterogeneity was located within the right ITR (**Fig 6d**).

### pAAV2ST: a stabilized rAAV cloning vector

We next tested whether the right ITR in the pAAVcis plasmid could be stabilized by increasing the spacing with respect to the origin of replication. A 1,797 bp DNA fragment containing the tetracycline resistance gene from pBR322 was inserted between the right ITR and the origin to produce the entry vector pAAV2ST (**Fig 7a**). To assess the stability of a construct that more closely resembles a typical rAAV designed for therapeutic gene transfer, we replaced the 1,569 bp NotI insert containing the GFP expression cassette with an all-in-one CRISPR/SaCas9 system previously developed by Feng Zhang and colleagues for the *in vivo* disruption of the murine *Pcsk9* locus (17). The total size of the resulting plasmid, pAAV2ST-Pcsk9sg1, was 8,722 bp (**Fig 7b**).

**Figure 7.**
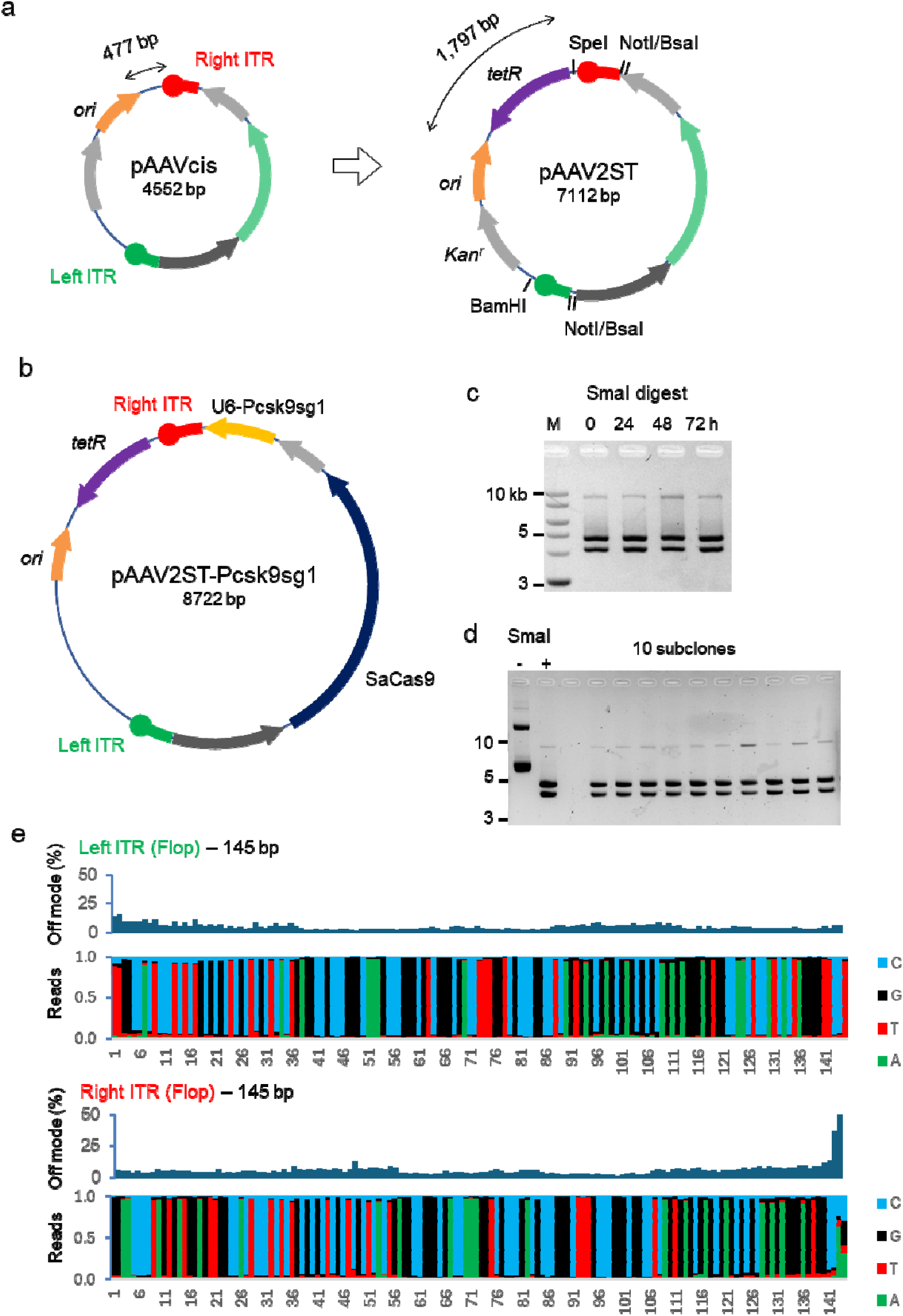
Stable propagation of ITRs flanking a 4,750 bp CRISPR/SaCas9 system. (a) The *tet^r^*selection marker from pBR322 was inserted between the adjacent *ori* and ITR elements in pAAVcis to generate the novel entry vector pAAV2ST (ST = stable). The *ori* and right ITR in pAAV2ST are 1,797 bp apart, as shown. The GFP cassette can be excised by digestion with NotI. Alternatively, digestion with BsaI facilitates directional cloning by the Golden Gate method (31). (b) The GFP expression cassette was replaced by an SaCas9 insert programmed to target the murine *Pcsk9* locus (17). (c) Bacteria harboring the pAAV2ST-Pcsk9sg1 plasmid were serially cultured for 72 h. DNAs extracted at 24-h intervals were digested with Smal. (d) pAAV2ST-Pcsk9sg1 DNA from a 10-mg gigaprep was left uncut or Smal-digested before gel electrophoresis, as indicated. Digested DNAs from 10 independent subclones were analyzed on the same gel. (e) Nanopore read pileup analysis of the ITRs from the large-scale pAAV2ST-Pcsk9sg1 DNA preparation. Nucleotide positions are shown on the horizontal axes.

The pAAV2ST-Pcsk9sg1 vector digested to near-completion with SmaI and appeared to be structurally stable during serial culture (**Fig 7c**). A sample of plasmid DNA, prepared at large scale for packaging, was predominantly supercoiled with no detectable linear species (**Fig 7d**). Subclones derived from this population exhibited low proportions of high molecular weight DNA after SmaI digest, as was previously observed in pBR-AAV2(-). Nanopore sequencing of this highly amplified preparation detected no mutated subpopulations. Both ITRs appeared to be homogeneous outside of a focal region at the 3’ end of the right ITR (**Fig 7e**).

The pAAV2ST-Pcsk9sg1 plasmid insert was packaged into AAV8 capsids by a commercial vendor using standard methods. Based on a qPCR-based quality control analysis by the vendor, a total of 5.61E+13 viral genomes were packaged, which was well in excess of the 1.0E+13 specification. Following purification, two main particle populations were resolved by mass photometry. A peak at 3,700 kDa representing empty capsids accounted for 29.5% of the population; 70.3% of the particles were full capsids, resolved in a 5,200 kDa peak (**Fig 8a**). We next visualized individual capsids in-house by negative staining followed by TEM (**Fig 8b**). This analysis indicated that 79.7 ±3.0% of the capsids were full and 20.2±3.0% were empty.

**Figure 8.**
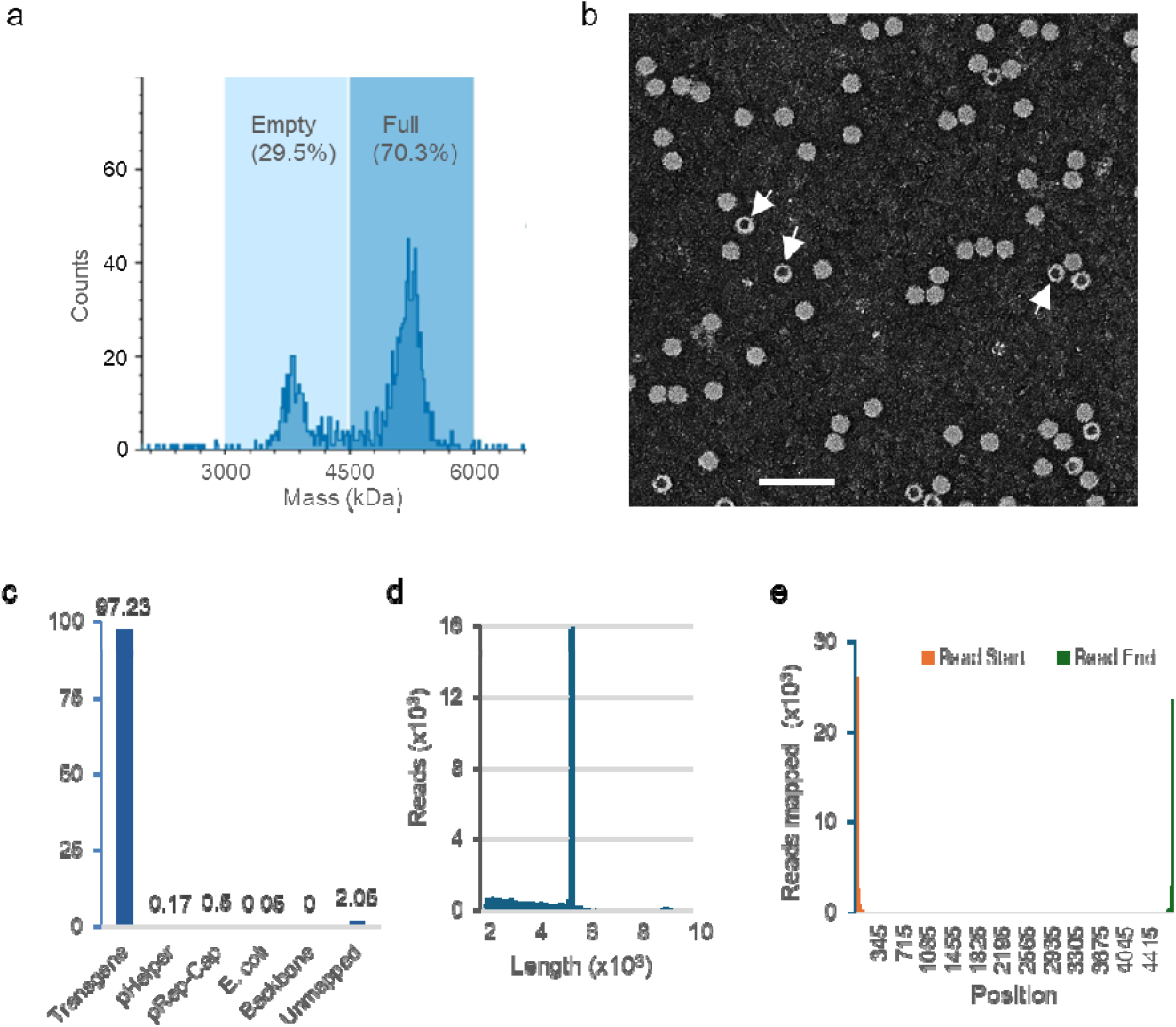
Analysis of a viral stock rescued from pAAV2ST-Pcsk9sg1. (a) Mass photometry analysis of gradient-purified AAV8 particles. (b) Purified virus particles visualized by transmission electron microscopy at 57,000x magnification. Examples of empty capsids are indicated with white arrows. Scale bar = 100 nm. (c) The distribution of the reads aligning to the transgene cargo, the Rep-Cap and helper plasmid DNAs, the *E. coli* genome, and unmapped reads are indicated. (d) Length distribution of encapsidated genomic DNAs, as assessed by nanopore DNA sequencing. (e) The start and end coordinates of all reads mapping to the ITR-flanked transgene.

The integrity of native encapsidated DNA was evaluated by direct nanopore sequencing, using a newly developed workflow. More than 98% of the resulting reads mapped to the transgene cassette (**Fig 8c**) and corresponded in size to the full length of the pAAVST-Pcsk9sg1 insert (**Fig 8d,e**).

### A widely disseminated ITR mutation

An important consequence of ITR instability is the clonal expansion of mutant subpopulations. An 11-bp ITR deletion (referred to hereafter as Δ11) was previously found in the common entry vector pAAV-MCS (16). A recent survey of sequence-validated rAAV plasmids in the Addgene repository by Radukic et al. (23) suggested that internally deleted ITRs are common. We sought to determine the prevalence of the specific Δ11 ITR mutant among all plasmids deposited with Addgene and asked whether this mutation showed a consistent positional relationship to the bacterial origin of replication.

A multi-stage bioinformatic screening pipeline was established to assess ITR sequences in the Addgene plasmid repository (**Fig 9a**). Sequence data were derived from a comprehensive public Addgene bulk download comprising approximately 1.34 Gb of plasmid sequence. A metadata-based screen identified 7,491 candidate AAV-related plasmids, which were then evaluated by sequence-level ITR detection. This analysis identified 7,041 plasmids with at least one detectable AAV ITR. Most sequence-verifiable AAV plasmids contained two inferred ITRs, with 6,824 plasmids classified as two-ITR vectors, 213 as one-ITR vectors, and 4 as three-ITR-like records. Of the total 13,873 ITR calls, 13,752 (99.1%) were classified as flop isomers and 121 (0.9%) were classified as flip.

**Figure 9.**
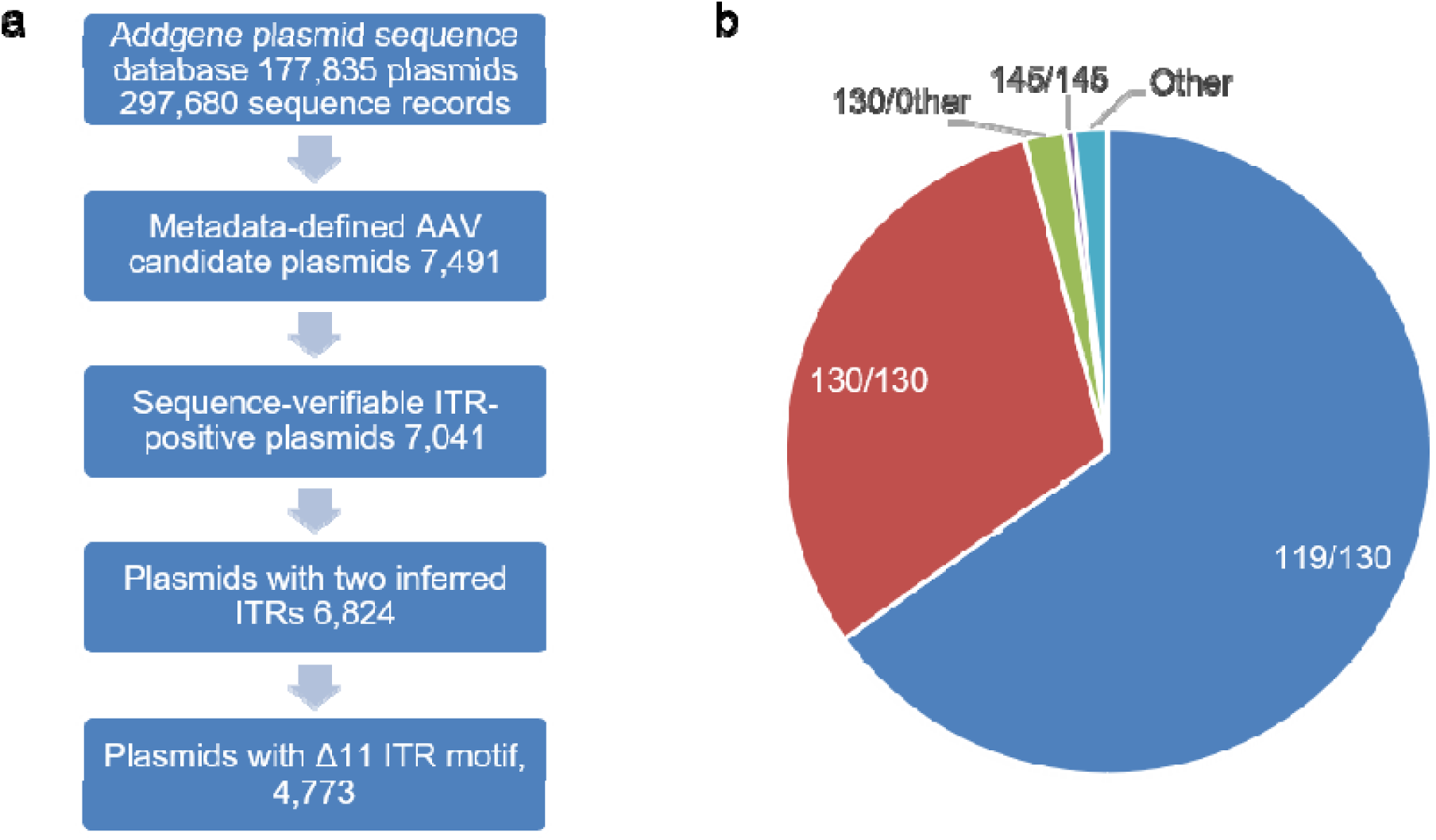
Sequence-based survey of ITR architectures in Addgene AAV *cis* plasmids. (a) Analysis pipeline The Addgene bulk plasmid sequence dataset contained 177,835 plasmids and 297,680 sequence records. Metadata filtering identified 7,491 candidate AAV plasmids, of which 7,041 contained sequence-verifiable AAV ITRs. Most ITR-containing plasmids contained two inferred ITRs. (b) Structural classes among 6,824 two-ITR plasmids. The two ITRs are denoted by their respective lengths, as first ITR/second ITR. The dominant architecture consisted of one 119-bp ITR with the Δ11 mutation and one 130-bp ITR. Approximately one-third of these plasmids contained two 130-bp ITRs. A much smaller number of cis plasmids featured two full-length 145-bp ITRs.

We next performed an independent exact-motif search for a 34-bp sequence corresponding to the Δ11 mutation. This search identified 4,773 plasmids with at least one exact Δ11 motif, representing 67.8% of ITR-positive plasmids and 69.8% of plasmids with two or more inferred ITRs. At the plasmid level, 4,763 plasmids contained a single exact Δ11 motif, whereas only 10 contained two exact Δ11 motifs. No plasmid containing an exact Δ11 motif lacked a sequence-detectable ITR, supporting the specificity of the motif-based search.

Because our experimental data suggested that ITR instability was influenced by proximity to the *ori* element, we asked whether the Δ11 ITRs were preferentially located near the annotated origin of replication. GenBank annotations were used to identify bacterial plasmid replication origins, and coordinate compatibility between the bulk Addgene sequences and GenBank files was confirmed in a test set. We restricted the origin-proximity analysis to plasmids with exactly two inferred ITRs and exactly one exact Δ11 motif in the representative full sequence. Of the 4,773 Δ11-positive plasmids, 25 were excluded because they did not meet these criteria or could not be analyzed. Among the remaining 4,748 plasmids, the Δ11-containing ITR was the ITR closest to the bacterial origin in 4,524 cases (95.3%) and was not closest to the origin in 224 cases (4.7%). Thus, across thousands of deposited AAV plasmid sequences, the Δ11 deletion is strongly enriched in the origin-proximal ITR, consistent with asymmetric, position-dependent ITR instability.

Interestingly, a small number of plasmids contained two exact Δ11 motifs. Manual inspection of these 10 cases indicated that 9 contained two separated ITR-associated Δ11 loci, whereas one appeared to reflect a local assembly or linearization-related duplication rather than two independent mutant ITRs.

In a majority of the AAV plasmids distributed by Addgene the ITR elements were either 130-bp or 119-bp in length (**Fig 9b**). The 130-bp ITRs presumably originated from the 15-bp truncation that was created during the construction of psub201. The additional Δ11 deletion in the *ori*-proximal ITR reduced the total size to 119 bp. This recurrent 11-bp ITR mutation is located within the C/C’ hairpin (**Fig 10a**), where it is predicted to disrupt base pairing that stabilizes the native T-shaped ITR conformation.

**Figure 10.**
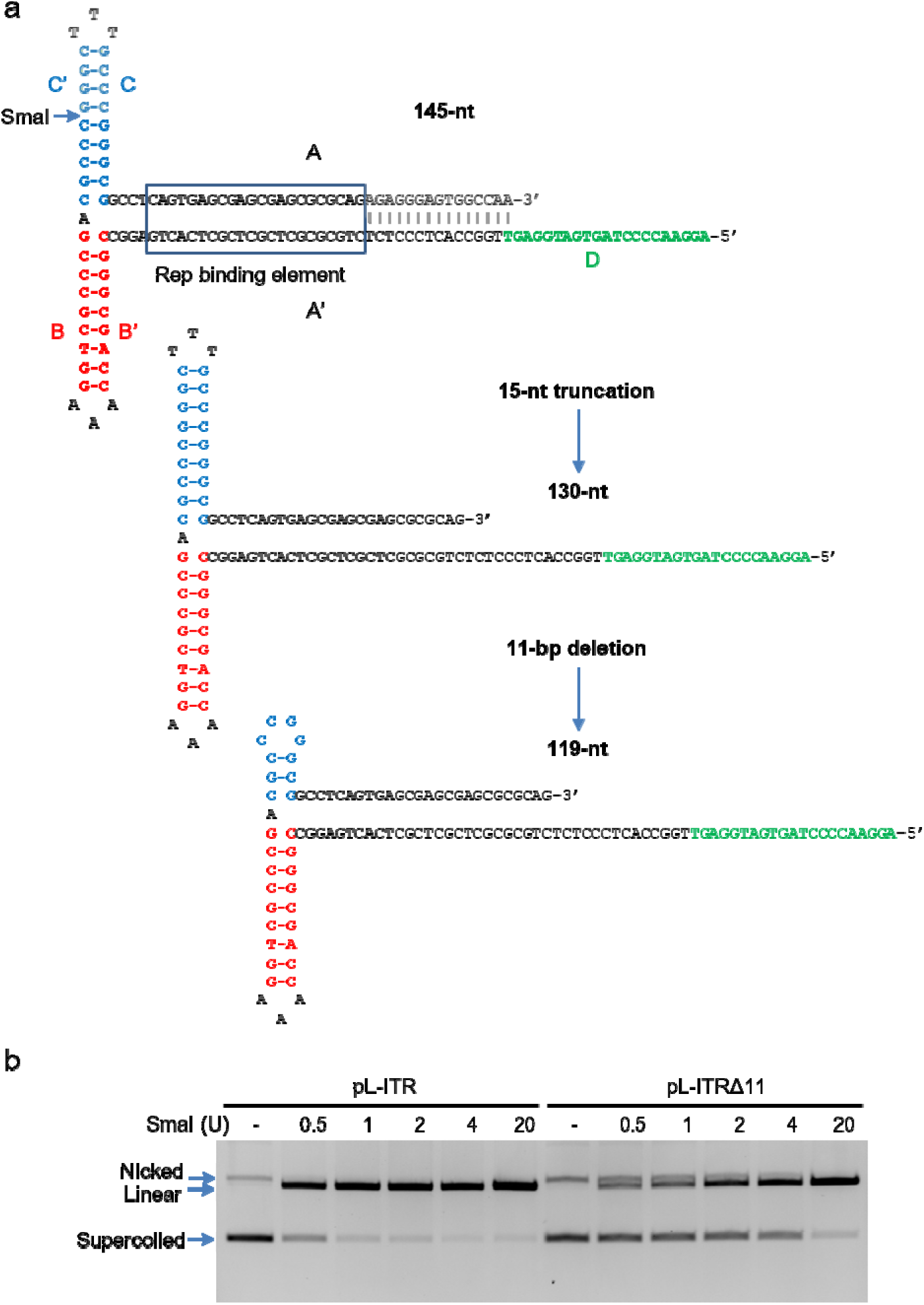
ITRs represented in published rAAV *cis* plasmids. (a) The native 145-nt ITR contains a duplex region (A/A’), two hairpins (B/B’ and C/C’), and a single-stranded domain (D). The binding element recognized by the Rep protein is boxed. Shaded portions represent the nucleotides eliminated by an 11-nt internal deletion and a 15-nt truncation commonly found in contemporary recombinant AAV vectors. The C/C’ duplex forms a duplex Smal recognition site, as indicated. (b) A 130-nt ITR was created by an ancestral 15-nt truncation. (c) A common 119-nt ITR contains the 15-bp truncation and the 11-nt internal deletion Δ11. (d) A 145-nt ITR plasmid (pL-ITR) and a variant containing the 11-bp internal deletion (pL-ITRΔ11) were synthesized individually. The two plasmids were digested with increasing amounts of SmaI and resolved on a native agarose gel. The positions of supercoiled uncut, linear, and nicked species are as indicated.

### Impact of mutant ITR sequences on genomic integration

We created two single-ITR plasmids, pL-ITR and pL-ITRΔ11, that respectively contained a 145-bp ITR and a 134-bp ITR with the 11-bp deletion, but which were otherwise identical. The common deletion eliminates one of the two SmaI sites present in the unmutated ITR. One consequence of the internal deletion is the predicted loss of duplex structure at the core of the SmaI recognition sequence, 5’-CCCGGG-3’ (**Fig 10a**). Since only the duplex form of the SmaI recognition site is a substrate for digestion, we were able to use SmaI to probe the structural equilibrium between the duplex and native forms of the mutant ITR (concept illustrated in **Fig 4a**.) The mutant ITR was more resistant to digestion at limiting concentrations of enzyme, suggesting a stable native conformation (**Fig 10b**). The wild-type ITR, which should be sensitive to SmaI whether in the native or duplex conformation, was digested nearly to completion at limiting amounts of enzyme, as expected.

To begin to explore whether this altered hairpin could impact the propensity for viral integration into the host genome, we devised a new assay for viral integration. This assay was designed to simultaneously measure non-homologous integration, which is largely dependent on micro-homologies shared by the ITR and integration sites, and homologous integration, which is dependent upon the homology-dependent repair of larger templates. We predicted that an increased propensity for insertional mutagenesis would be reflected in a decreased ratio of homologous integration/non-homologous integration.

We modified an existing gene targeting vector, pSEPT-p53, that integrates into the human *TP53* locus with high efficiency (24). The pSEPT-p53 construct features a synthetic exon promoter trap that permits the selectable marker to be expressed only when it is integrated downstream of an endogenous promoter, thus enriching for clones that arise from homologous integration events (**Fig 11a**). The homology arms in pSEPT-p53 were designed so that the neomycin resistance gene (neo^r^) would replace an 80-bp segment in the first coding exon. The cassette itself lacks a promoter, so episomal DNAs cannot confer drug resistance; non-homologous integration events can only be detected if *neo^r^* is positioned downstream of an active promoter.

**Figure 11.**
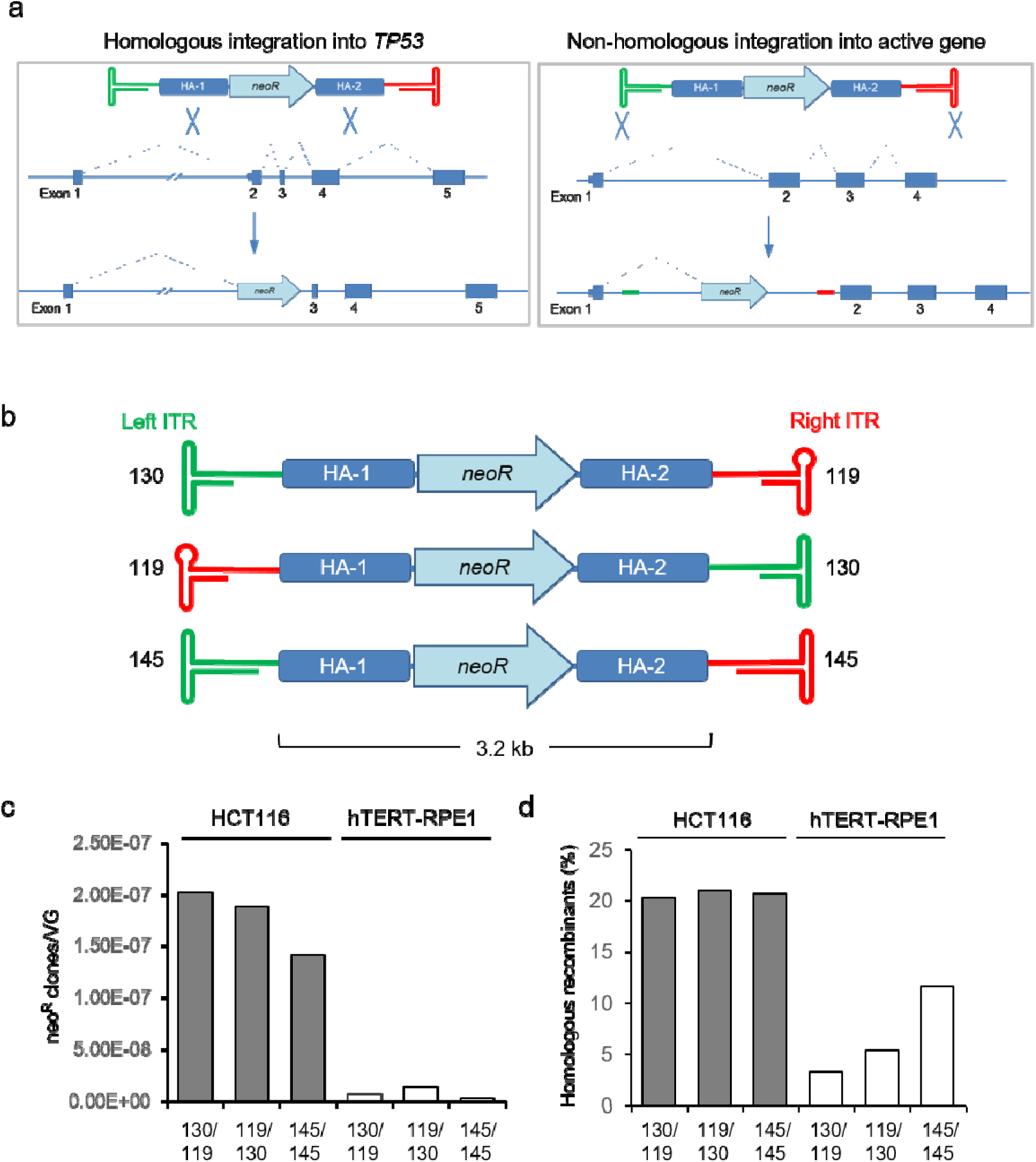
Assessing relative levels of homologous and non-homologous integration. (a) The pAAV-MCS derivative pSEPT-p53 is a promoter-trap vector designed for homologous integration of viral genomes into the human *TP53* locus (18). Two homology domains (HA-1 and HA-2) define a small deletion in exon 2, which is replaced by a promoterless *neo^r^* cassette upon homologous integration (*left*). This construct also integrates into off-target sites that share microhomology with ITRs; ITR remnants should be present at the junctions between the endogenous gene and the targeting construct (*right*). The relative risk of insertional mutagenesis can be inferred from the ratio of targeted to non-targeted integration events. (b) The NotI-NotI insert from pSEPT-p53 was cloned in both orientations into the asymmetric pAAV-MCS backbone and into pAAVcis. The lengths in bp of the right and left ITRs, with respect to the insert orientation, are indicated. (c) The number of neomycin-resistant (neo^r^) colonies derived from each of the three vectors was scored three weeks after infection of HCT116 cells (gray bars) or hTERT-RPE1 cells (white bars). The vectors are denoted as left ITR length/right ITR length across the horizontal axes. (d) The proportion of neo^r^ colonies that resulted from homologous integration into the *TP53* locus.

Originally constructed in the pAAV-MCS backbone, the pSEPT-p53 plasmid has a 130-bp ITR upstream of the selectable marker and a downstream 119-bp ITR (**Fig 11b**). The insert was flanked by NotI sites, so we were able to flip the insert and thereby create a comparable vector in the opposite orientation. The NotI-NotI gene-targeting insert was also introduced into the pAAVcis plasmid in place of the GFP expression cassette (**Fig 7a**).

The integration of packaged viral genomes, as assessed by colony formation, was approximately 10-fold greater in the mismatch repair-deficient colorectal cancer cell line HCT116 than in the telomerase-immortalized human cell line hTERT-RPE1 (**Fig 11c**). In both cell lines, the virus derived from the construct with wild-type ITRs generated a smaller number of drug-resistant colonies than either of the two viruses produced from the ITR-mutant constructs. Whereas the proportion of homologous integrants in HCT116 was similar between the three viruses, the wild-type ITR construct was associated with a higher rate of homologous integration in hTERT-RPE1 (**Fig 11d**). In hTERT-RPE1, the virus derived from the construct with the 119-bp ITR in the upstream configuration appeared to integrate somewhat more frequently overall than the vector in the reverse orientation (**Fig 11c**) and generated a modestly higher proportion of homologous recombinants (**Fig 11d**).

## Discussion

The ITR elements from AAV2 have a well-known tendency to undergo deletion during repetitive rounds of plasmid amplification (12). This distinctive form of genetic instability leads to heterogeneity within plasmid populations and is a potential source of variability among virus preparations. In this study, we demonstrated that instability is not an intrinsic property of ITRs, but rather an idiosyncratic defect that stems from their positioning. By reconfiguring the founding AAV2 isolate pSM620 so that both ITRs were >1.4 kb from the pMB1 origin we were able to reduce ITR loss to a nearly undetectable level. A synthetic high-copy entry vector could be similarly stabilized by increasing the origin-to-ITR spacing. ITR integrity was maintained even when plasmids were propagated in generic cloning bacteria (*E. coli* DH5α) grown under standard culture conditions.

Long-read nanopore sequencing and per-position pileup analysis of raw reads provided an opportunity to quantify plasmid heterogeneity and thereby examine the positional effect at single-molecule resolution. We found that both ITRs were maintained during single-colony amplification of pSM620 (**Fig 3a,b**). However, extended culture under saturating conditions led to the deterioration of the left ITR and expansion of ITR-deleted subclones (**Fig 5a-d**). The asymmetric instability of ITRs in pSM620, which became apparent following extended culture, was mirrored in the synthetic construct pAAVcis (**Fig 6a-d**), supporting the conclusion that origin proximity was the determining factor.

The pMB1 *ori* functions in a directional manner but appeared to affect the stability of an adjacent ITR in either orientation. The asymmetric distribution of the flip/flop signal between the two ITRs of pAAVcis, and its absence in the origin-proximal left ITR of the low-copy pBR322-derived pSM620, suggests that replication fork geometry may be an additional contributing factor to the observed positional effects. In ColE1-type plasmids, a single unidirectional fork emanates from the origin. In pAAVcis, the origin is oriented so that the replication fork encounters the right ITR from its outer palindromic end, with the D-sequence replicated last (**Fig 6a**). Under this geometry, the sense strand of the ITR serves as the lagging strand template and can conceivably fold into the canonical T-shaped hairpin in which the flip/flop inner loop is exposed as a single-stranded loop.

While our study did not directly address the mechanistic basis of the observed positional effect, we hypothesize that the juxtaposition of the *ori* and an ITR creates RNA-DNA hybrids that may facilitate recombination. The pMB1 *ori* is a finely tuned RNA–DNA switch that is mediated by structural interactions (22). The origin itself is A/T-rich, favoring DNA unwinding. Plasmid copy number is determined by the interplay of two non-coding RNAs. A transcript called RNA II is transcribed from a start site approximately 550 bp upstream, forming a structure known as the R-loop that serves as the primer for DNA replication. An antisense RNA, RNA I, forms three hairpins that disrupt the R-loop, thereby exerting a suppressive effect on priming. In low-copy plasmids, the plasmid-encoded protein Rop stabilizes the interaction between the two RNAs and thus further enforces copy number control. Ultimately, the plasmid copy number is a function of the relative levels of these two RNAs and the stability of their interaction. Further study is needed to pinpoint which region of the complex *ori* structure may have directly impacted the conformational state of the ITR.

A comprehensive analysis by Radukic *et al.* (23) recently identified several factors that contribute to ITR instability, including origin proximity. One notable finding was that stalled DNA replication at ITR sequences triggered degradation by the SbcCD nuclease complex. Reportedly, the very close proximity of the *ori* and ITR elements (<300 bp) was a particularly strong determinant of ITR degradation. Culturing ITR-containing plasmids in SbcC-deficient strains at an elevated temperature of 42°C reportedly resulted in significant improvements in ITR stability. Radukic *et al.* noted that ITRs were stable at a minimum distance of 438 bp from the plasmid origin. In contrast, under the experimental conditions of our study, we observed that stability could be improved by increasing the ITR-to-*ori* spacing from 474 bp to >1 kb. Further investigation would be needed to determine whether the improved methods established by Radukic *et al.* might synergize with increased ITR-*ori* spacing.

In many instances, the consequences of ITR mutation can be mitigated during genome replication and packaging. Small alterations are repaired by recombination between the two ITRs (15), while nucleotides missing from either end of the AAV genome, such as the terminal 15 bp that were eliminated in psub201 (19), are restored by template-directed DNA synthesis. Accordingly, small deletions affecting a single ITR have been experimentally demonstrated to have little impact on virus production (14). Nonetheless, the continued prevalence of a single 11-bp deletion among published vectors (16,23,25) and in a majority of sequence-verifiable AAV plasmids in the Addgene repository (**Fig 9a**) clearly demonstrates the limitations of self-repair mechanisms.

Large ITR deletions are more obviously problematic. Virions can be efficiently rescued from single-ITR plasmid vectors, but the encapsidated genomes tend to be incomplete and contaminated with snapback genomes (11). Simple measures, such as the use of vectors that are optimized for ITR stability, would be a straightforward and low-cost approach to minimizing ITR loss and thereby limiting these types of defects.

The stabilization of the plasmid ITR elements did not eliminate the production of empty capsids (**Fig 8a,b**). Nonetheless, the AAV preparation produced from the pAAV2ST-Pcsk9sg1 stock was of high quality and titer. Based on our TEM analysis (**Fig 8b**), the proportion of full AAV8 capsids produced from the pAAV2ST-Pcsk9sg1 vector was very similar to the full capsid fraction (78.1%) reported by Khaparde *et al.* (26), who described the production of a research-scale AAV8 stock purified by ultracentrifugation and analyzed by TEM.

The impact of vector stability on the overall quality of the rescued virus is difficult to quantify. Our efforts to reduce the likelihood of complete ITR loss in plasmid DNA destined for packaging did not appear to compromise packaging efficiency, but it is not clear whether there was any significant improvement in the standard metrics of quality. It is important to acknowledge that the pAAV2ST-Pcsk9sg1 stock was packaged and purified once at a commercial facility. Additional study would be required to confirm that viruses produced from the pAAV2ST entry vector are of consistently high quality and yield.

The remarkable persistence of rAAV-mediated transgene expression is one of the important attributes of these vectors. Most intracellular viral genomes stabilize in a double-stranded episomal state, but a small proportion integrate into the host genome (27,28). These integration events, while relatively rare, have raised the concern that rAAV-mediated genomic mutagenesis could increase the risk of cancer (29).

The genomic integration of wild-type AAV into specific sites in the human genome requires the viral proteins Rep68 and Rep78, which bind to a recognition sequence within the ITR (**Fig 10A**) and introduce a nick within a motif called the terminal resolution site (24). This nicking reaction initiates strand transfer between the ITR and sites in the host genome that share a short region of homology. rAAVs lack the Rep gene and therefore do not appreciably integrate into these sequence-defined loci. However, rAAV integration occurs at low frequency, generally through non-homologous end joining mechanisms involving the ITR hairpins. Non-homologous integration appears to be non-random, as rAAV genomes disproportionately integrate at numerous “hot spots”, which are commonly found within CpG islands and actively transcribed genes.(28) The single-stranded region of the rAAV genome can provide an excellent substrate for homologous recombination, and indeed, such vectors have been used extensively for gene targeting (25).

Our genomic integration assay exploited the high efficiency with which a previously developed rAAV gene targeting vector could disrupt *TP53* (18). We reasoned that an elevated rate of non-homologous integration should result in a larger number of integration events overall, and a commensurate decrease in the rate of homologous recombination. In both cell lines, the vectors that retained the 11-bp deletion stably integrated at a higher rate compared with the vector that had two complete ITRs (**Fig 11C**). Only in the hTERT-RPE1 cells was this higher integration frequency associated with a lower rate of homologous integration. The elevated rate of homologous recombination in HCT116 is largely attributable to defective mismatch repair (MMR), which eliminates the dependency on complete sequence identify between the vector and target site (30). It is possible that a higher stringency for homologous recombination in MMR-proficient hTERT-RPE1 cells accentuated the differences between the vectors. These cells may therefore be ideally suited for these types of analyses. Overall, the rAAVs integrated into HCT116 cells at a significantly higher rate. This finding may simply reflect a general tendency of cancer cells to tolerate genetic alterations, but we cannot rule out more idiosyncratic differences between these two cell lines, such as their AAV infectivity.

The integration data presented here should be interpreted with caution. Our goal was to explore a ratiometric approach to assessing viral integration. While our study suggested that mutant ITRs have measurable effects on the balance of non-homologous and homologous integration, the design of this preliminary experiment precluded rigorous statistical analysis. Additional experimentation would be needed to confirm these findings with multiple viral stocks and replicate infections. From a mechanistic standpoint, it would be important to identify the specific junctions between the host genome sequences and the integrated vectors to evaluate whether the ITR mutation might have influenced the integration site biases. Nonetheless, the assay described here represents a novel approach to integration analysis that could be more broadly applied. Additional refinements to this method could improve it considerably. For example, the use of a vector that efficiently targets a selectable gene like HPRT would make for a simpler readout of homologous integration. Considering that rAAV vectors harboring the 11-bp deletion have very likely been used in clinical trials, we believe that additional study would be well justified.

This study had several other limitations. During the reconfiguration of the pSM620 plasmid we eliminated the G/C homopolymeric tracts that flank the AAV2 genomic insert. A remnant of the original cloning process, these sequences have previously been shown to increase the overall instability of the ITRs (12). However, the rescue of instability in the synthetic construct pAAVcis, which lacks the homopolymeric flanking sequences in pSM620, suggests that origin proximity is the more significant destabilizing factor.

Our reliance on several different marker genes is another potential limitation. The pBR322 vector confers resistance to both ampicillin/carbenicillin and tetracycline/doxycycline (**Fig 4a**). The positioning of the AAV2 inserts disrupted one or the other of these markers. As a result, pSM620 had to be propagated in the presence of doxycycline, while carbenicillin selection was required for maintenance of the pBR-AAV2 plasmids. We cannot rule out distinct selective pressures as a contributing factor to the observed differences between the low-copy plasmids. However, the stabilization of pAAV2ST involved kanamycin selection only, again suggesting that origin proximity was paramount.

## Acknowledgements

The pX602-AAV-TBG::NLS-SaCas9-NLS-HA-OLLAS-bGHpA;U6::BsaI-sgRNA plasmid was kindly provided by Feng Zhang (Addgene plasmid 61593). The study was supported by the Foundation of the National Institutes of Health (FNIH) on behalf of the Accelerating Medicines Partnership Program Bespoke Gene Therapy Consortium (AMP BGTC).

## Author contributions

Conceptualization: V.J-R., S.L., F.B.; Investigation: S.L., A.P., F.B.; Writing-Original Draft Preparation: V.J-R.,F.B.; Writing-Review & Editing: V.J-R., S.L., A.P., F.B.; Project Administration: F.B..

## Notes

https://github.com/fredbunz-lgtm/aav-itr-pileup

https://www.ncbi.nlm.nih.gov/sra/?term=PRJNA1439403

### Competing Interest Statement

The authors have declared no competing interest.

